# Is *Oculudentavis* a bird or even archosaur?

**DOI:** 10.1101/2020.03.16.993949

**Authors:** Zhiheng Li, Wei Wang, Han Hu, Min Wang, Hongyu Yi, Jing Lu

## Abstract

Recent finding of a fossil – *Oculudentavis khaungraae* Xing et al. 2020, entombed in a Late Cretaceous amber – was claimed to represent a humming bird-sized dinosaur^1^. Regardless of the intriguing evolutionary hypotheses about the bauplan of Mesozoic dinosaurs (including birds) posited therein, this enigmatic animal demonstrates various morphologies resembling lizards. If *Oculudentavis* was a bird, it challenges several fundamental morphological differences between Lepidosauria and Archosauria. Here we reanalyze the original computed tomography scan data of *Oculudentavis*. Morphological evidences demonstrated here highly contradict the avian or even archosaurian phylogenetic placement of *Oculudentavis*. In contrast, our analysis revealed multiple synapomorphies of the Squamata in this taxon, including pleurodont marginal teeth and an open infratemporal fenestra, which suggests a squamate rather than avian or dinosaurian affinity of *Oculudentavis*.

---

Instead of demonstrating synapomorphies of the Aves, *Oculudentavis* show multiple characters that have never been found in any previously known birds or non-avian dinosaurs. One of the most bizarre characters is the absence of an antorbital fenestra (Fig. 1a, b). Xing et al.^1^ argued the antorbital fenestra fused with the orbit, but they reported the lacrimal is present at the anterior margin of the orbit^1^. This contradicts the definition of the lacrimal in all archosaur including birds since lacrimal always forms the caudal margin of the antorbital fenestra^2^. In addition, closure of the antorbital fenestra is rare in archosaurs^3-5^, and most birds and all the known Cretaceous birds do have a separate antorbital fenestra^6^.

**Figure 1.**
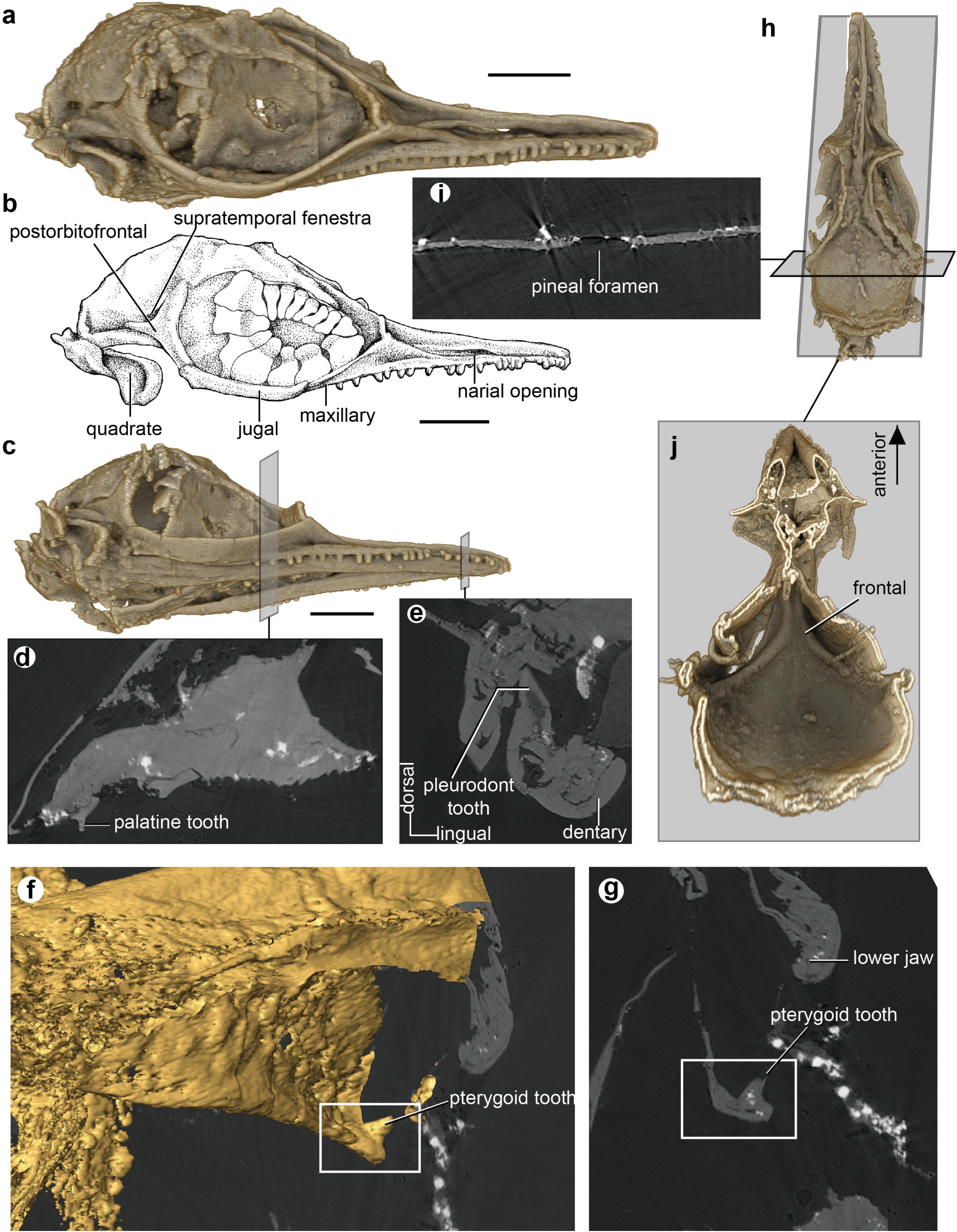
Reanalysis of the cranial anatomy of *Oculudentavis khaungraae* Xing et al. 2020^1^ (holotype, HPG-15-3) based on the original computed tomography (CT) scan data. **a**, Three-dimensional CT reconstruction of the skull in right lateral view. **b**, Line drawing of the skull in right lateral view, showing the absence of quadratojugal in *Oculudentavis*. **c**, Skull in ventrolateral view. **d** and **e**, Two-dimensional CT slices through the palatine (**d**, showing a palatine tooth) and the dentary (**e**, showing a typical pleurodont tooth). **f** and **g**, Pterygoid tooth shown in three-dimensional reconstruction of the skull (**f**) and in a coronal plane through of the skull (**g**). **h**, Skull in dorsal view. **i**, A coronal CT slice through the skull roof showing the pineal foramen. **j**, Skull in ventral view, with the lower jaw and palate removed to show the ventral surface of the frontal. **a**-**c**, scale bar: 2 mm; **d**-**j**, not to scale.

Another highly questionable feature in *Oculudentavis* is the maxilla extending caudally to the level of mid-orbit and forming half of the ventral margin of the orbit (Fig. 1b), which is extremely unusual in Aves. In most crown birds, the maxilla terminates anterior to the orbit. The ventral margin of the orbit is formed by the jugal^2,7^. This is also the condition among Mesozoic birds, including *Archaeopteryx*^8,9^, *Sapeornis*^10^, enantiornithines^6^ and ornithuromorphs^6^. In *Ichthyornis*, maxilla is elongate and extends further caudally beneath the jugal^11^ which means the ventral margin of the orbit is still mostly composed by the jugal, different from *Oculudentavis*. In addition, we need to note that the skull of *Jeholornis* was incorrectly reconstructed with a maxilla extending most of the orbit, followed by a shortened jugal^1^, which present a mislead similarity between the skull of *Oculudentavis* and *Jeholornis*. However, the maxilla of *Jeholornis* is short and most of the ventral margin of the orbit is formed by the elongate jugal followed by the quadratojugal^6^, in stark contrast with *Oculudentavis*.

In *Oculudentavis*, the maxillary tooth row extends caudally to the rostral half of the orbital. Among most Mesozoic birds, maxillary tooth row ends well cranially to the cranial margin of the orbit^6^. In contrast, at least four teeth are located beneath the ventral margin of the orbital, and the last one even ends below the rostral third point of the orbit in *Oculudentavis*.

Although Xing et al. mentioned that the scleral ring and dentition of *Oculudentavis* resemble lizards^1^, they failed to recognize that pleurodont dentition is diagnostic for squamates^12^. The maxillary and dentary teeth are ankylosed to the jaw with their labial side (Fig. 1e), and replacement teeth develop posterolingual to the functional teeth. The authors also stated that the tooth implantation appears to be acrodont to pleurodont. However, there is no evidence for acrodonty based on our reexamination of the original CT scan data.

In comparison, dinosaurs have thecodont teeth that develop in tooth sockets, with replacement teeth developing beneath the functional teeth. Although the Late Cretaceous ornithuromorph bird *Hesperornis* retain teeth in a groove (tooth sockets fused together)^13^, it is clearly distinguishable from the pleurodont dentition in *Oculudentavis*. Non-archosaurian dentition of *Oculudentavis* has also been interpreted as the result of miniaturization^1^. To our best knowledge, there is no concrete evidence suggesting such a drastically change of dentition in miniaturized archosaurs. Pleurodont dentition falsifies the dinosaurian or even archosaurian affinity of *Oculudentavis* — instead it supports the squamate affinity of this new species.

Another unambiguous squamate synapomorphy in *Oculudentavis* is the loss of the lower temporal bar. In the original publication^1^, a complete orbit was illustrated on the left side of the skull with an unnamed piece of bone between the jugal and postorbitofrontal^1^. In addition, the anterior margin of the quadrate articulates with an unlabeled bone. The misleading illustration suggests that the quadratojugal might be present in *Oculudentavis*. On the basis of the original CT scan data, we demonstrate that the orbit on the left side of the skull is crushed. The left jugal is not preserved. The right side of the skull preserves a complete orbital region, which shows the jugal has a smooth posterior margin, lacking contact with the quadrate. The quadratojugal is absent (Fig. 1a and b), which means the infratemporal fenestra is open in *Oculudentavis* – a condition shared with all squamates but not dinosaurs or birds^12,14^.

Additional morphologies of *Oculudentavis* that contradict its avian affinity include the presence of the parietal foramen (Fig. 1i), the separate ventral down growths of frontal (Fig.1j), as well as palatal teeth present on palatine and pterygoid (Figs. 1d, k, and l)

*Oculudentavis* means “eye-tooth bird”, yet neither the eyes (scleral ring) nor the teeth suggest this new species was a bird. Xing et al^1^ assigned this enigmatic animal to Aves based on superficial appearances, such as the exterior contour of the dome-shaped cranium and slender rostrum^1^. However, all the extended discussions, including the morphological changes related to miniaturization and the ocular morphology, lost their foundation with a problematic phylogenetic placement of this animal. In addition, multiple unambiguous characters support the squamate affinity of *Oculudentavis*, including the loss of quadratojugal, pleurodont marginal teeth, and presence of palatal teeth (Figs. 1 and 2). The original phylogenetic analysis by Xing et al. suffers from biased sampling of taxa^1^. Our new morphological discoveries suggest that lepidosaurs should be included in the phylogenetic analysis of *Oculudentavis*.

**Figure 2.**
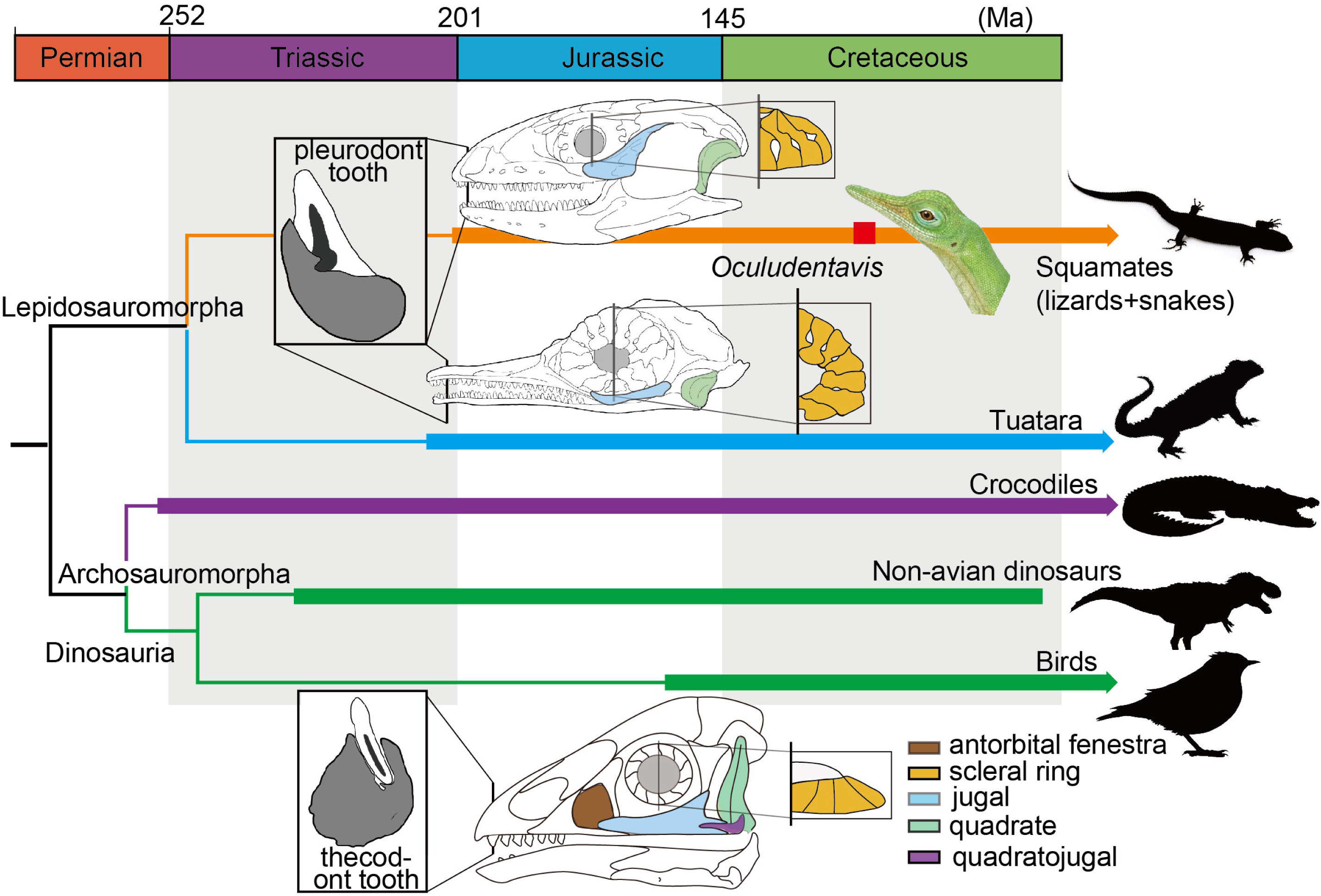
Simplified reptile family tree, illustrative drawings showing the comparison of the skull in *Oculudentavis*, squamate (green lizard *Lacerta bilineata*) and bird (Cretaceous bird *Sapeornis*).

## Supporting information

Combined and re-rendered 3D format files in Drishti

## Methods and Data availability

The original CT scan data was obtained upon request from the authors of original paper^1^. Two 3D format files (9.5G in total) were combined into one and re-rendered in Drishti 2.6.6 (https://github.com/nci/drishti/releases)^15^. Scan data were analyzed in Avizo (www.thermofisher.com) and imaged in Adobe photoshop (www.adobe.com). For more scanning, 3D reconstruction and data information see ref 1.

## Acknowledgement

We thank G.L. for providing the original CT scan data for reanalyzing. We thank colleagues at Institute of Vertebrate Paleontology and Paleoanthropology, Chinese Academy of Sciences for their great help during preparing and writing the manuscript.

## Author Contributions

All authors designed the project, analyzed and discussed the data, and wrote the manuscript. All authors contributed equally. Correspondence and requests for materials should be addressed to M.W. or to H.Y.Y. or to J.L.

